# The number of Follicle Stem Cells in a Drosophila ovariole

**DOI:** 10.1101/2021.10.25.465475

**Authors:** Daniel Kalderon, David Melamed, Amy Reilein

## Abstract

A paper by Reilein et al (2017) presented several fundamental new insights into the behavior of adult Follicle Stem Cells (FSCs) in the Drosophila ovary, including evidence that each ovariole hosts a large number of FSCs (14-16) maintained by population asymmetry (Reilein et al., 2017), rather than just two FSCs, dividing with largely individually asymmetric outcomes, as originally proposed (Margolis and Spradling, 1995; Nystul and Spradling, 2007). Fadiga and Nystul (2019) contest some of these conclusions on the basis of their repetition of a multicolor lineage strategy used by Reilein et al (2017) and repetition of earlier single-color lineage analysis. Here we outline a number of shortcomings in the execution and interpretation of those experiments that, in our opinion, undermine their conclusions. The central issue of general relevance concerns the importance of comprehensively analyzing all stem cell lineages, independent of any pre-conceptions, in order to identify all constituents and capture heterogeneous behaviors.

## Introduction

Each type of adult stem cell can best be defined as the group of cells that maintain production of at least one different cell type throughout adult life (Clevers and Watt, 2018; Post and Clevers, 2019). Lineage analyses are extensively used to identify and characterize stem cells but generally involve examining the products of single cells over limited periods of time (Blanpain and Simons, 2013; Fox et al., 2008). To make deductions about a stem cell community from such single cell data it is necessary to examine many stem cell lineages comprehensively and without bias, allowing variant behaviors to be captured at appropriate frequencies, and to do so over a variety of time periods in order to capture lineages of all lifetimes.

Drosophila oogenesis affords the possibility of good cell visualization, efficient methods for genetic lineage labeling and a very straightforward temporal method for identifying lineages originating in the stem cells upstream of Follicle Cells (FCs). The initial definition of FSCs exploited these advantages and represented a big step forward, dictating dogma over the next two decades (Margolis and Spradling, 1995). Over a period of time, culminating in the publication of Reilein et al (2017), our group realized that earlier studies had effectively made the assumption that FSCs were maintained by invariant single cell asymmetry. Once this assumption was discarded in favor of a comprehensive analysis of rigorously defined FSC lineages, supplemented by comprehensive three-dimensional and temporal analyses, it became clear that FSCs are maintained instead by population asymmetry, that there are many more FSCs, of varied lifetimes, than previously thought and that FSCs in fact also maintain a second ovarian cell type-non-dividing Escort Cells (ECs)(Reilein et al., 2017).

Fadiga and Nystul (2019) recently contested some of these conclusions. Here we outline why we believe the conclusions of Reilein et al (2017) are sound, fully answering the challenges raised by Fadiga and Nystul (2019).

## Results and Discussion

Drosophila females produce eggs throughout adult life. Eggs develop from a sequence of increasingly large egg chambers that first bud from the germarium at the anterior end of each ovariole (Figure 1) (Duhart et al., 2017). An egg chamber contains germline cells surrounded by a monolayer of Follicle Cells, with specialized Polar Cells at either end and Stalk Cells connecting neighboring egg chambers. Here and in Reilein et al (2017) we refer to all of these cells, as well as somatic cells associated with stage 2b and 3 germarial cysts as Follicle Cells (FCs). The defining feature of an FC is that it associates with a germline cyst, moves out of the germarium with that cyst and continues that association as the egg chamber matures. The somatic cells that are not themselves FCs but maintain production of FCs are called FSCs. This defining difference is determined experimentally by timing (Figure 1). A marked FC produces a lineage associated with a single cyst that passes through an ovariole within five days (Margolis and Spradling, 1995). Any lineage mark, initiated at a defined time by heat-shock induction of a relatively short-lived “Flp” recombinase, that labels FCs anywhere in an ovariole beyond this 5d time window necessarily originated in an FSC, and the lineage is referred to as an FSC lineage. This definition is consistent with the first definition of FSCs (Margolis and Spradling, 1995) and with a general principle that can be applied to all adult stem cells (Clevers and Watt, 2018; Post and Clevers, 2019).

**Figure 1.**
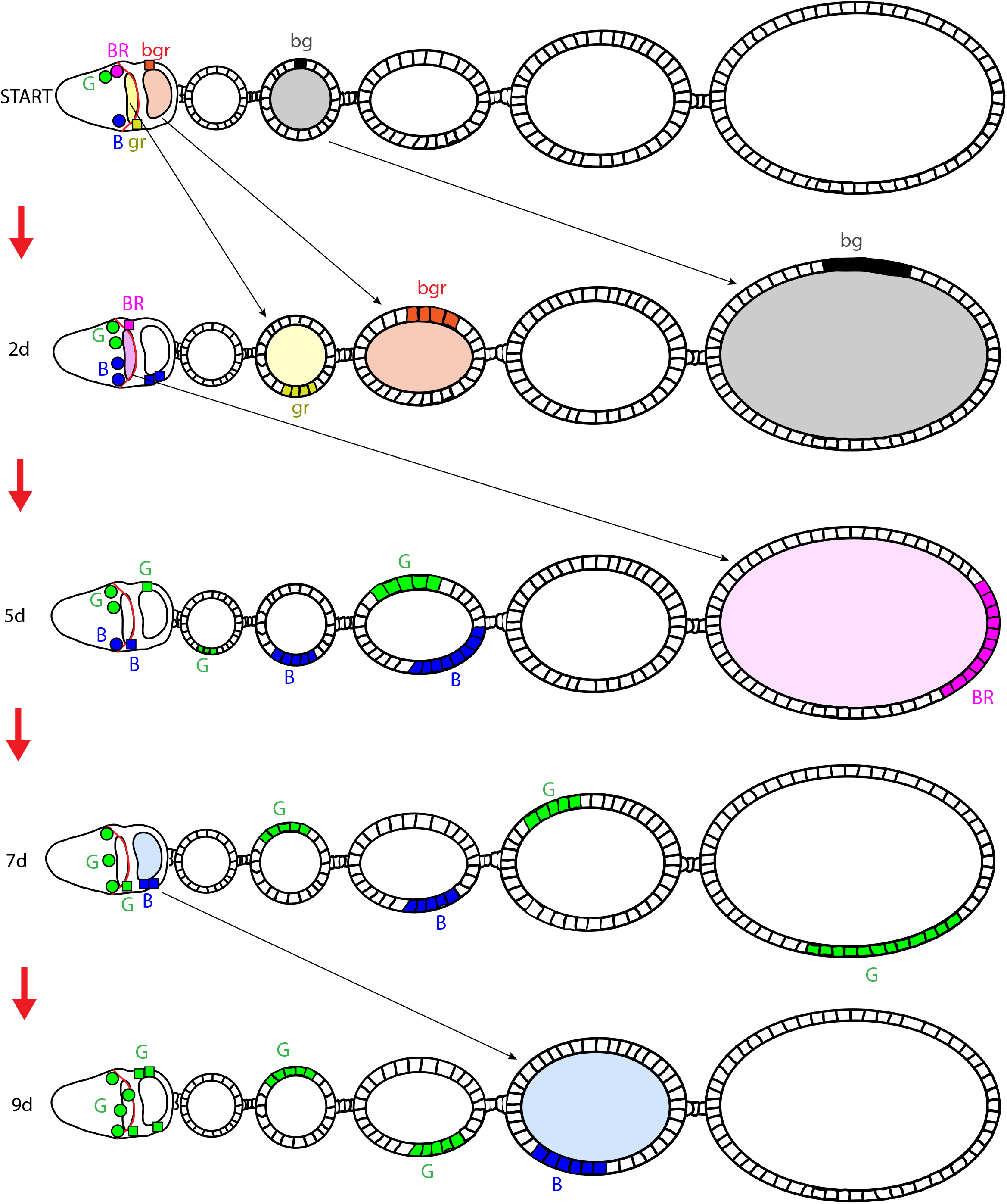
Temporal definition of a marked lineage initiated in a FSC. The cartoon illustrates the germarium (left), housing a lens-shaped stage 2b germline cyst (yellow), followed by a more spherical stage 3 germline cyst (orange) and five egg chambers of increasing size. Shortly after heat-shock (by about 0.5d and labeled “START”) a subset of proliferating FSCs and Follicle Cells (FCs) have undergone recombination and are depicted by six examples in different colors. By 2d, after about 3 cycles of egg chamber budding, all initially labeled cells associated with a germline cyst (yellow, orange and black squares) have amplified into a contiguous patch and have moved posteriorly (to the right) together with their associated germline cysts, which are shown with matching colors. The characteristic of stable association with a germline cyst defines a FC. By 5d, even the earliest FC on a stage 2b cyst at START (yellow; gr), together with its entire lineage, has exited the ovariole (which extends beyond the five egg chambers shown). Any labeled FCs present anywhere in the ovariole five or more days after heat-shock induced recombination must therefore derive from a recombination event in a cell upstream of an FC; all such cells are defined as FSCs (circles with upper case labels). The behavior of individual FSCs is the subject of FSC lineage tracing experiments. Three possible examples are illustrated. The magenta (BR; blue plus red) FSC survives for a couple of 12h cycles of egg chamber budding but it is then lost, becoming a FC on a stage 2b cyst by 2d. The lineage originating in the magenta (BR) FSC is still present as a single FC patch at 5d but it is lost from the ovariole by 9d. The blue (B) FSC survives longer, occasionally producing FCs (blue squares) but at some time between 5d and 7d the last blue FSC is lost. The last blue FC produced (at 6.5d) matures into a FC patch in the fourth egg chamber at 9d, with no other remnant of the blue lineage. The green (G) FSC initially produced no FCs but amplified and then green FSCs continued to produce FCs intermittently, while maintaining an FSC population that varies in number over time, with four green FSCs present at 9d. In the example illustrated, the green lineage would be scored as a FSC lineage with at least one surviving FSC. The blue lineage would be scored at 9d as an FSC lineage with no surviving FSC (and it can be inferred that there was a surviving blue FSC at 5d). Note also that the blue FSC lineage would only be detected if at least four egg chambers were scored. The purple lineage did originate in an FSC but it would be missed because the ovariole is scored at 9d (to leave no possibility of lineages originating from an FC) rather than after the minimal possible time of 5d. The experimental definition of a FSC lineage does not rely on knowing or assuming where FSCs are located. The exact dynamics of somatic cell associations with cysts in the germarium are not known but are also not relevant for defining an FSC lineage. If a cell initially associated with a cyst in the germarium but did not emerge with that cyst in a budded egg chamber, the lineage derived from that cell would, appropriately, be scored as originating in an FSC. Additional studies, based on FSC lineages with only single candidate FSCs are required to define the locations of FSCs; those studies place the majority of FSCs immediately anterior to the anterior border of strong Fas3 staining (red line at the posterior edge of stage 2b cysts) or one cell further anterior.

### Multicolor lineage Analysis

Reilein et al. (2017) devised a genetic system that allowed Flp-induced recombination at *FRT* sites near the base of two chromosome arms (2L and 2R) to allow the generation of independent lineages with five distinguishable recombinant genotypes (B, G, BG, BR and GR where B, G and R indicate the presence of *lacZ*, GFP and RFP markers, respectively). This system was used to generate FSC lineages with as much color diversity as possible in order to count the number of distinguishable FSC lineages in each ovariole. We reported the presence of an average of 4.5 differently colored FSC lineages over 50 ovarioles 9d after marking (Reilein et al., 2017). We repeated this analysis at various times after FSC labeling to reveal that individual FSCs are frequently lost, especially soon after marking, and to estimate that each ovariole included about 16 FSC lineages prior to any losses.

Fadiga and Nystul used our donated reagents to conduct analogous experiments. First, they reported fewer distinguishable FSC lineages (only 2.1 on average at 9d). The underlying explanation is simple. Fadiga and Nystul did not induce recombinant genotypes at a high frequency (Figure 2B, C). Under these circumstances most FSC lineages have the same color and the approach cannot count how many FSC lineages are present. Fadiga and Nystul also did not detect all FSC lineages present because they scored ovarioles only up to the second or third egg chamber (see Figure 1). Consequently, the basic requirements of high efficiency labeling and detection were not met and the number of stem cells was substantially underestimated.

**Figure 2.**
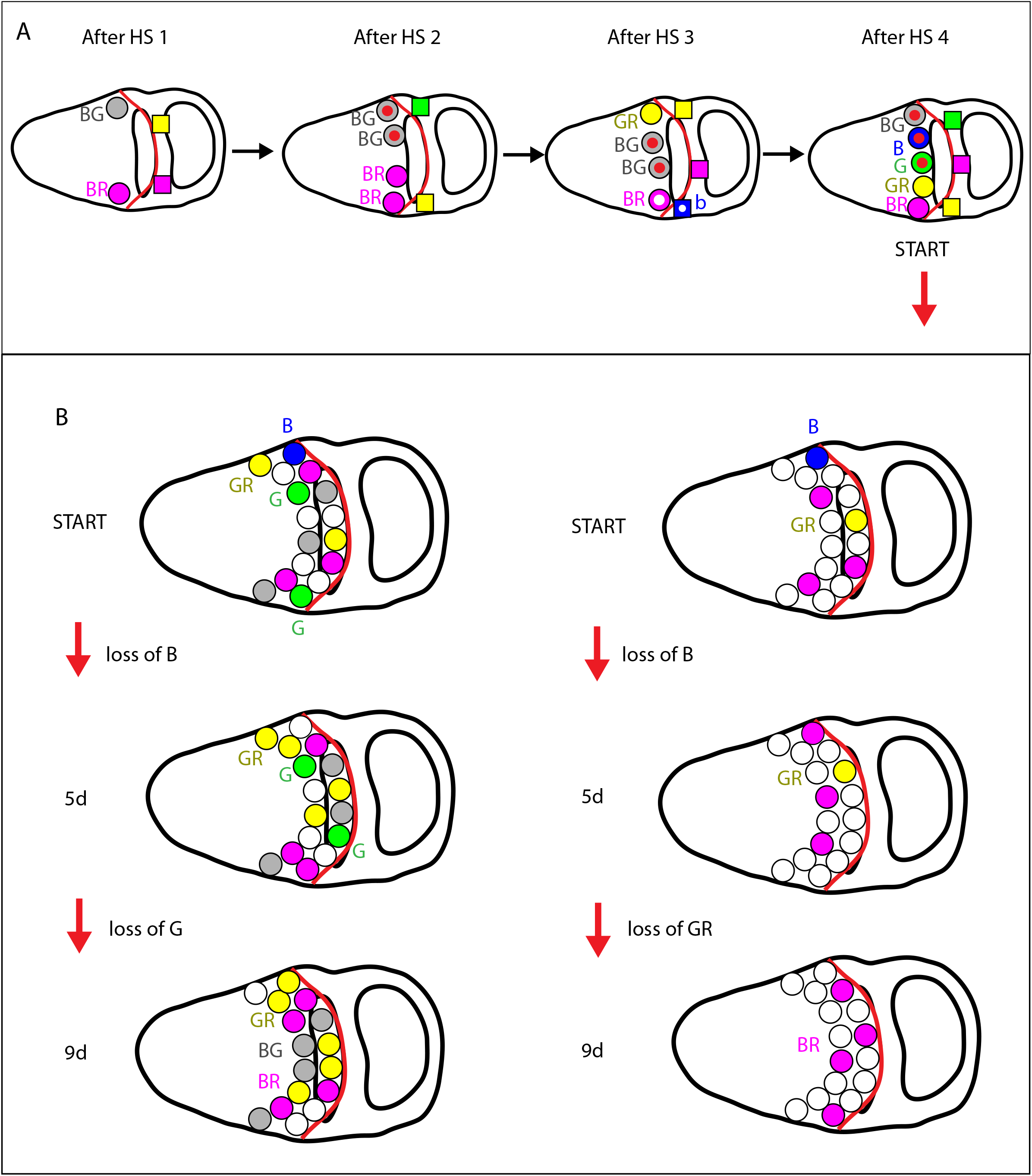
Multiple heat-shocks and the key role of high frequency marking of FSC lineages in multicolor lineage analysis for counting the number of FSC lineages. (A) The pictured germaria illustrate a possible sequence of events when using four spaced heat-shocks (“HS”) to elicit four rounds of possible recombination events. As in Fig. 1, FSCs are shown as circles and the anterior border of strong Fas3 staining (red) is shown, close to the posterior surface of the stage 2b germline cyst. A grey (BG) FSC generated by recombination shortly after the first heat-shock later duplicates and then one BG FSC undergoes recombination after the fourth and final heat-shock to produce B and G daughter FSCs (“subclones”, indicated by a red dot). No further recombination takes place because there are no further heat-shocks and the timed experiment begins now (“START”), with ovarioles examined 9d later. Those ovarioles will show the presence of B, G and BG FSCs (among others) if those FSC lineages are not lost early. It will be deduced correctly that the B, G and BG lineages derived from an FSC of the corresponding color at the start of the experiment. That is equally true whether B and G are “subclones”, as shown here, or if they had no lineal relationship to each other or the BG FSC, as for the GR (yellow) and BR (magenta) FSCs. Whether the cell has a red dot or not (indicating its past history) has no influence on how it will behave or its identity as an FSC. All FCs with recombinant genotypes will not be represented in ovarioles 9d later, regardless of whether any are “subclones” (white dots) or not. (B) The pictured germaria illustrate typical plausible scenarios when recombination occurs at high frequency (left) or low frequency (right) among FSCs. As outlined in the Recombination Frequency section of Multicolor lineage analysis in the archive deposition xxx, the data of Reilein et al (2017) and Fadiga and Nystul (2019) conform roughly to frequencies of producing recombinant colors of 0.7 and 0.3 per cell, respectively. For p=0.7 (left) there will, on average, be 11 recombinant colors among the 16 pictured FSCs, with lower frequencies of RFP-negative colors (because RFP/+ and RFP/RFP are both RFP-positive). A frequency of recombinant colors of 0.3 (right) will, on average, produce 5 recombinant genotypes among the 16 FSCs, with a very low frequency of B and G genotypes, which can only be generated by two recombination events. It is therefore expected that all five recombinant FSC colors will usually be represented among an average of 11 recombinant FSCs for p= 0.7 (the parental color is the sixth and will also generally be present) and usually no more than three different recombinant colors (B, BR and GR are shown) will be represented among an average of five recombinant FSCs for p= 0.3. Moreover, each of the three colors will necessarily be represented by only a small number of cells (here B and GR have just one FSC). Over time, colors represented initially by one or a small number of FSCs are the most likely to disappear. Reilein et al (2017) deduced that about 2/3 of initial FSC lineages are lost over 9d. The illustrated examples conform to these expectations. By 5d the initial single blue (B) FSC is lost in both scenarios (left and right). Between 5d and 9d, for p= 0.7 (left) the green (G) lineages, initially represented by 2 FSCs are lost, whereas yellow (GR) lineages, also initiated by two FSCs, are retained. For p= 0.3 (right) the yellow (GR) lineage, initiated by only one FSC (fewer than in the germarium on the left), is lost between 5d and 9d. In both cases the number of FSCs in other lineages changes stochastically (due to FSC division and FSC loss), with the same behavior pictured for the three magenta (BR) FSCs that were present initially in both cases. In 9d samples, in the ovariole derived from high frequency FSC labeling (left) there will be four colors of FSC lineages with surviving FSCs (BG, BR, GR and the parental white lineage) and another FSC lineage color without a surviving FSC (G; scored also as a surviving FSC at 5d). For the ovariole derived from low frequency FSC labeling there are only two FSC lineage colors with surviving FSCs (BR and parental) and one additional FSC lineage color without a surviving FSC (GR), which would be missed by Fadiga and Nystul if the FSC was lost prior to 7d because ovarioles were not scored beyond 2-3 egg chambers (illustrated in Fig. 1). These depicted scenarios and outcomes are commensurate with the observations of Reilein et al (2017) that more than 50% of ovarioles had five or more colors of FSC lineages, and more than 40% had four or more colors of FSC lineages with surviving FSCs at 9d, whereas Fadiga and Nystul (2019) reported that most ovarioles have fewer than three FSC lineages. The cartoons illustrate that both sets of results are consistent with the presence of multiple FSCs (sixteen here) but that, crucially, the number of FSCs present can only be measured accurately if different FSCs are made visible by labeling most, or all of them with different colors and detecting the presence of differently colored lineage by examining the whole length of the ovariole.

Second, concerns were raised about the methods used by Reilein et al (2017). Fadiga and Nystul suggested that we assumed 100% recombination at both *FRT* locations in all cells to derive FSC numbers from counts of differently colored lineages. We did not. Our data showed an average recombination frequency of about 2/3 at each *FRT* and we used that observation to infer the frequency of lineages that shared the same color and hence estimate the number of lineages present. Fadiga and Nystul also suggested that the use of multiple heat-shocks is inappropriate because “subclones” may be generated. The number of heat-shocks is of no conceptual importance (Figure 2). During the series of heat-shocks a diversity of genotypes is generated with any manner of clonal relationships among the final set of FSCs. After the final heat-shock there are no further genotype changes and the timed experiment that defines each FSC lineage begins.

Finally, Fadiga and Nystul cite a high frequency of recombinant lineages in the absence of deliberate heat-shock and a staining pattern that should not occur (loss of both *lacZ* and GFP at an undisclosed frequency) in their experiments to allege that the multicolor genetic system is somehow flawed. Neither problem was present in the studies of Reilein et al (2017) or in a recent repeat we conducted.

### Single-color lineage analyses

The number of FC-supplying FSCs can also be estimated by measuring the proportion of all FCs contributed by a single FSC. It is, however, crucial to ensure that only one FSC is labeled in each ovariole and to score every ovariole in which an FSC was originally marked to capture the potentially heterogeneous behavior of individual FSCs. Indeed, FSC lineages examined at a single time point show a great variety of appearances, with varied and sporadic contributions of marked FCs (Figures 3 and 4) (Reilein et al., 2017). Fadiga and Nystul (2019) reported a very similar distribution of FSC lineage phenotypes. However, they measured FC contributions only for the small proportion of ovarioles where marked FCs are present in the germarium and the first two egg chambers. This introduced a large bias, leading to a much-inflated estimation of about 50% of FC being marked in each lineage. Reilein et al (2017) included all FSC lineages with marked FCs and at least one surviving FSC to find an average FC contribution per lineage of 12% (multicolor) or 15% (MARCM) (Figure 3 and 4). Those lineages included an average of 3.3 FSCs (and hence a variable number between about 1 and 3.3 during the course of the experiment), so that the best estimation of FC production per FSC is closer to half of those values (6-8%, consistent with 14-16 FSCs).

**Figure 3.**
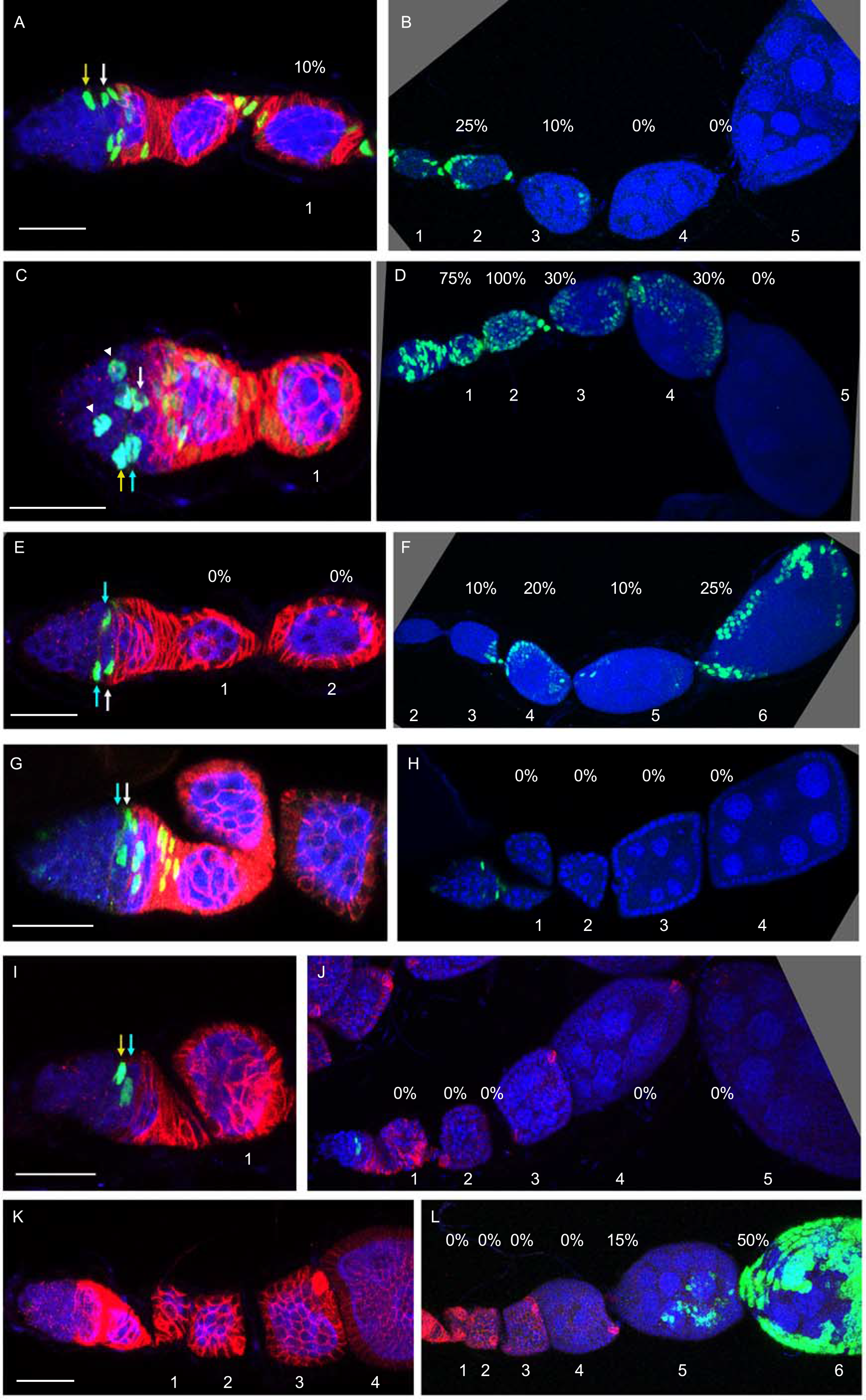
MARCM FSC lineages illustrate the heterogeneity of FSC behaviors, frequent FSC loss and FSC amplification. (A-L) A selection of MARCM clones from a single experiment examining ovarioles 9d after clone induction. Panels on the left include the germarium and early egg chambers, while panels on the right show the remainder of the same ovarioles; egg chambers are numbered to make clear overlapping regions. GFP-labeled cells (green) in each lineage are shown together with DAPI staining of all nuclei (blue) and Fas3 antibody staining (red) in select panels. Each image includes only a limited set of z-sections, so not all GFP-positive cells are visible. FSCs are indicated by arrow colors as being in layer 1 (white), layer 2 (blue) or layer 3 (yellow). The estimated proportion of the FC epithelium of each egg chamber that has GFP label is shown by the indicated percentage values (calculated by looking through all z-stacks). The final average value of 15% labeled FCs was calculated from all lineages (over sixty) with at least one marked FSC and FC patch (Reilein et al., 2017). (A-D) Ovarioles with marked FSCs and FCs in both the germarium and one or more egg chambers (36/148 ovarioles; 120/186 egg chambers with marked FCs; average of 4.4 FSCs). (E, F) Ovariole with marked FSCs and FCs in one or more egg chamber but not in the germarium (16/148 ovarioles; 50/84 egg chambers with marked FCs; average of 2.1 FSCs). (G, H) Ovariole with marked FSCs and FCs in the germarium but not in any egg chamber (10/148 ovarioles; average of 4.4 FSCs). (I, J) Ovariole with marked FSCs but no FCs (21/148 ovarioles; average of 1.7 FSCs). (K, L) Ovariole with marked FCs but no FSC (14/148 ovarioles). Several of the lineage categories above also included ECs. Remaining ovarioles (not shown) included ECs but no FSC or FC (20/148) or no marked cells (31/148). Only 24 lineages included marked FCs in the germarium and both of the first two egg chambers (the criterion used by Fadiga and Nystul for selecting lineages to score FC contributions) out of a total of 117 ovarioles with marked cells, of which 76 are definitive FSC lineages judged by the presence of marked FCs.

**Figure 4.**
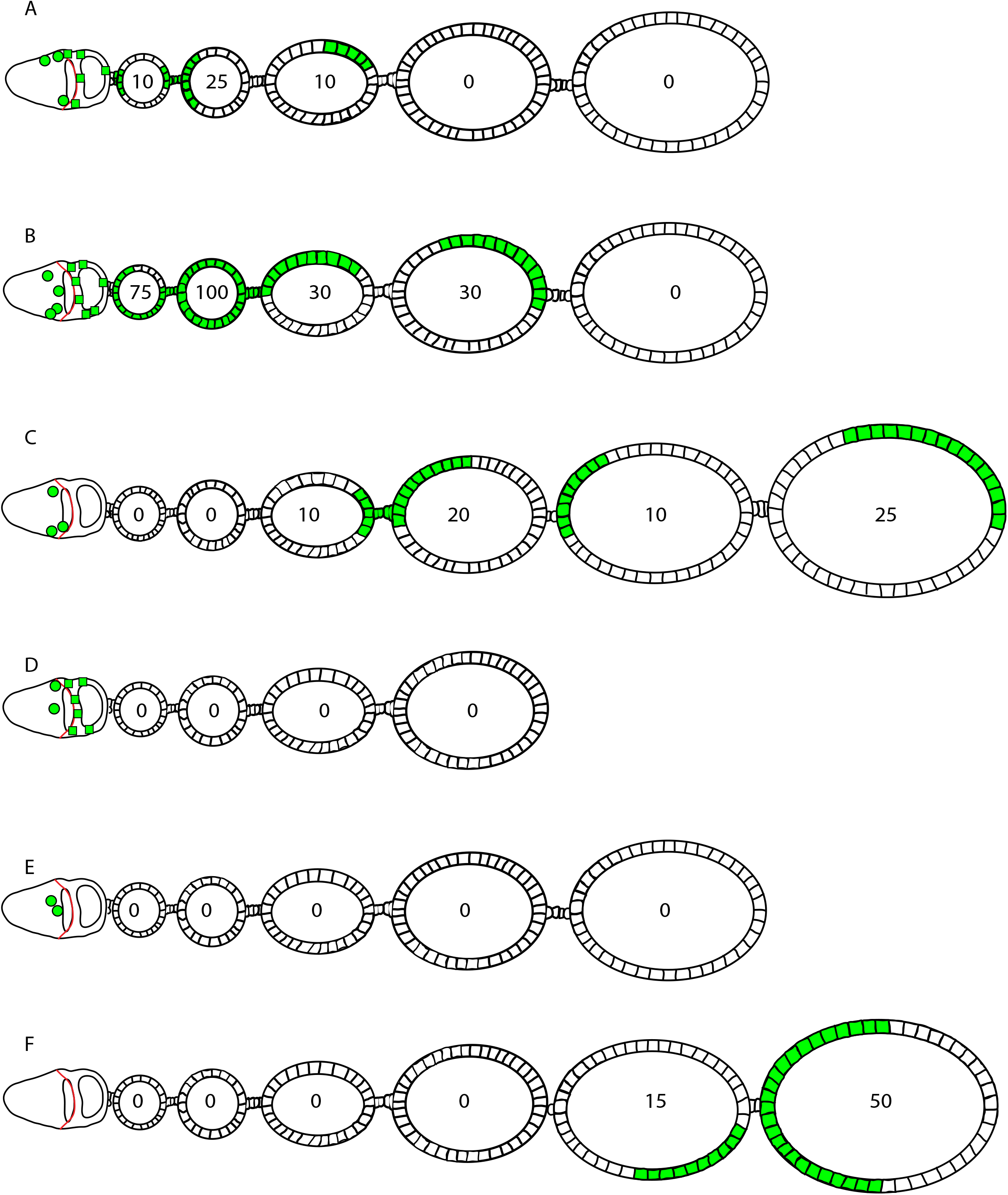
Summary of MARCM FSC lineages illustrated in Figure 3. (A-F) Diagrammatic representation of the types of FSC lineage shown in Figure 3. As explained in more detail in the legend to Figure 3, the relative frequency of the different patterns among all ovarioles examined was (A, B) 24% in sum, (C) 11%, (D) 7%, (E) 14% and (F) 9%.

The variability in FSC lineage appearances has several sources. FSCs divide at different rates, contribute FCs stochastically, and the number of FSCs per lineage changes over time because individual FSCs are lost or amplify at a high rate (Reilein et al. (2017)). Hence, it is imperative not to exclude any FSC lineages (as Fadiga and Nystul did) and it is inevitable that samples analyzed at 7-9d will contain many lineages maintained by multiple FSCs (while extinct lineages with zero FC contributions cannot be recognized and counted), requiring that the number of FSCs be counted in order to deduce the contribution of marked FCs per FSC.

The same issue explains earlier mis-conceptions of FSC stability. Early studies regarding FSC stability measured the loss of FSC lineages over periods subsequent to an initial 7d starting point (Margolis and Spradling, 1995; Nystul and Spradling, 2007); the rate of loss greatly over-estimated FSC stability because lineage loss required the loss of many FSCs, not just a single FSC.

The study of Fadiga and Nystul, and other studies preceding Reilein et al (2017) did not acknowledge the possibilities that FSCs may behave heterogeneously with high rates of loss and amplification of individual FSCs. Instead, the appearance of a subset of FSC lineages with continuous FC contributions was originally labeled as prototypical, effectively embodying an assumption that each FSC is independently maintained by a series of repeated divisions with an asymmetric outcome (Margolis and Spradling, 1995). Thereafter, this assumption was often treated, inappropriately, as a defining feature, leading to exclusion of the majority of FSC lineages from analyses. The latter lineages and their prevalence is equally evident in the reports of Reilein et al (2017) and Fadiga and Nystul (2019); those lineages definitively derive from FSCs, according to the fundamental and unambiguous definition of FSCs by temporal lineage criteria, as outlined earlier (Figure 1).

Reilein et al (2017) identified the location of FSCs by examining lineages in which candidate FSCs were found in only one location. FSCs were located throughout the circumference, adjacent to the germarium wall, and in two major rings or layers along the anterior-posterior axis, constituting a total of 14-16 cells. Almost identical results were derived from multicolor clones, with the advantage of illuminating the relative locations of FSCs of different colors, and MARCM clones, which most clearly identify all germarial cells (Reilein et al., 2017). Fadiga and Nystul (2019) did not identify FSC location using the critical criterion of examining clones with only a single candidate FSC or comprehensively score the locations of all candidate FSCs but instead assumed that the FSC will be the most anterior cell in a lineage. That approach cannot define FSC location.

In summary, the single-color analyses of Fadiga and Nystul (2019) provide inaccurate estimates of FSC number principally because they exclude the large fraction of FSC lineages that contribute few FCs, while their multicolor experiments did not induce and detect differently colored FSC clones efficiently enough to illuminate nearly all FSCs present. Reilein et al (2017) assayed FSC lineages comprehensively without bias, using three different approaches to reach a consistent conclusion of 14-16 FSCs per germarium.

## Acknowledgments

Research on this topic was supported by GM079351 awarded to DK. We thank Rachel Misner for comments.

## Supplementary Materials

A detailed critique of the central papers discussed in this necessarily abbreviated Scientific Correspondence to explain more carefully some of the assertions made and to include a detailed response to every issue raised by Fadiga and Nystul (2019).

